# Inferring When to Act from Temporal Regularities

**DOI:** 10.64898/2026.08.24.746547

**Authors:** Matteo Girondini, Greta Madonna, Tommaso Bertoni, Alberto Gallace

## Abstract

Adaptive behaviour often requires deciding when to act in the absence of an explicit sensory cue. While temporal expectations are known to optimize behaviour when anticipated events trigger responses, it remains unclear how learned temporal regularities are transformed into internally generated decisions. Here, we developed the Temporal Inference Task, a novel virtual reality paradigm designed to isolate this transformation. Participants repeatedly observed an identical visual sequence in which a virtual object approached their hand. On most trials, a brief tactile No-Go signal instructed them to withhold their response, whereas on the remaining trials its omission required them to respond. Because Go trials were never accompanied by an explicit cue, participants had to infer when the expected No-Go signal could no longer occur before initiating a response. Across blocks, the timing of the No-Go signal was systematically varied, allowing participants to learn distinct temporal regularities while Go trials remained physically identical. Participants systematically shifted their response timing according to the learned temporal regularities. This behavioural adaptation was accompanied by corresponding shifts in parietal alpha- and beta-band desynchronization, with steeper pre-response desynchronization consistently preceding faster responses. Together, these findings identify a candidate neural mechanism through which learned temporal regularities are transformed into internally generated decisions about when to act.

## Introduction

Temporal regularities enable the brain to anticipate future events by predicting when they are likely to occur, thereby optimizing behavior^1,2^. Previous work has shown that temporal predictions improve performance (e.g., reaction times, RTs) and modulate alpha- and beta-band oscillatory activity, particularly when actions are triggered by anticipated sensory events^3,4,5,6,7,8^. However, most studies have investigated temporal expectation within stimulus–response paradigms, in which the occurrence of an external sensory event directly triggers behavior. Yet, many real-world actions are not elicited by an explicit sensory cue but generated from an internal model of the temporal structure of the environment^9^. For example, when entering a revolving door, people continuously predict when the next compartment will reach the appropriate position and initiate their movement accordingly, rather than waiting for an explicit signal to step forward. Because the rotational speed can vary across revolving doors, successful behavior depends on continuously updating these expectations to determine the appropriate moment to initiate movement. Such situations require the brain to transform learned temporal regularities into internally generated decisions about when to act, rather than simply reacting to external events. How humans transform learned temporal regularities into internally generated decisions remains largely unknown. Here, we investigated the neural mechanisms through which learned temporal regularities give rise to the process of determining when to act in the absence of an explicit sensory trigger.

## Results

To address this question, we developed the Temporal Inference Task, a virtual reality (VR) paradigm designed to investigate how temporal expectations shape internally generated decisions. Participants (n = 25) repeatedly observed an identical visual sequence in which a virtual sphere approached the hand. On most trials (80%), a brief tactile stimulus instructed them to withhold their response (No-Go), whereas on the remaining trials (20%), its omission required a response (Go). Because Go trials were never accompanied by an explicit cue, participants had to infer when the expected No-Go stimulus could no longer occur before initiating a response. To manipulate temporal expectations, No-Go events occurred at a fixed time within each block: 300 ms before sphere–hand contact in the Early condition and 200 ms after sphere disappearance in the Late condition (Figure 1A). As Go trials were physically identical across conditions, any differences reflected the learned temporal distribution of No-Go events.

**Figure 1.**
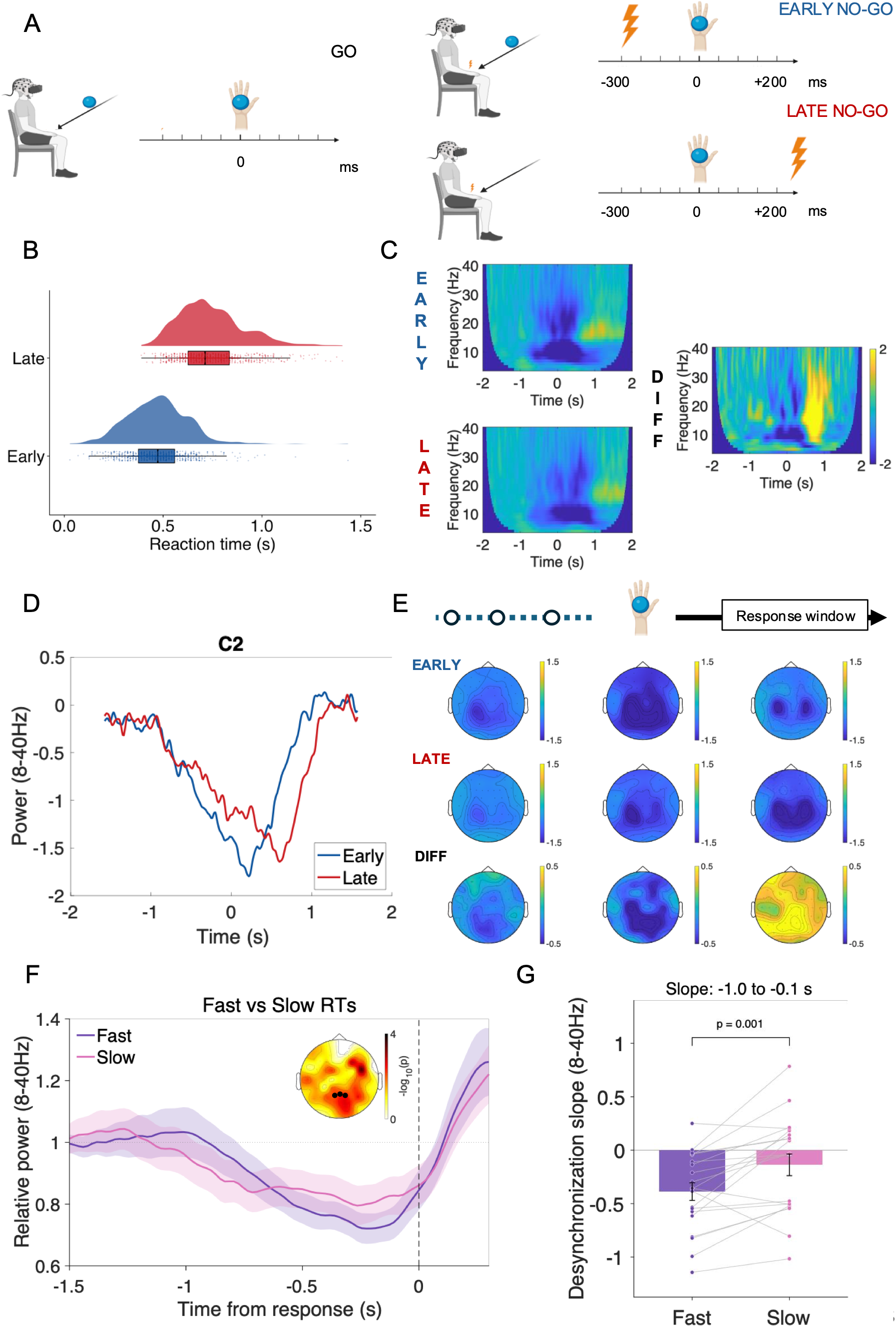
Learned temporal regularities shape inferential decisions through sensorimotor oscillatory dynamics. **(A) Temporal Inference Task.** Participants observed a sphere approaching their hand in virtual reality. On No-Go trials, a tactile stimulus instructed them to withhold their response; on Go trials, its omission required a button press. Across blocks, No-Go events occurred either before sphere–hand contact (Early; −300 ms) or after contact (Late; +200 ms), requiring participants to infer when it was safe to respond from the learned temporal distribution of No-Go events. **(B) Reaction time distributions in the Early and Late blocks**. Participants responded systematically earlier when No-Go events were concentrated before contact and later when they were concentrated after contact, demonstrating that response timing adapted to the learned temporal regularities of the environment. **(C) Time–frequency representations at electrode C2** for the Early and Late conditions and their difference (Early − Late). **(D) Time course of alpha/beta power (8–40 Hz) at electrode C2**. Oscillatory activity was temporally advanced in the Early relative to the Late condition, closely paralleling the behavioral shift in response timing. **(E) Topographical distribution of alpha/beta activity at representative time points**. Upper row: Early condition; middle row: Late condition; lower row: Early − Late difference. The difference maps reveal a centro-parietal predominance, consistent with the engagement of sensorimotor and parietal networks during temporally guided action preparation. **(F) Response-locked alpha/beta desynchronization**. Grand-average alpha/beta power (8–40 Hz), averaged across parietal electrodes (Pz, P1, P2), aligned to the button press. Faster responses were preceded by a steeper decrease in oscillatory power than slower responses, indicating a more rapid build-up of sensorimotor desynchronization before action initiation. **(G) Pre-response desynchronization slope**. Mean slope of alpha/beta desynchronization computed over the response window (−1.0 to −0.1 s before the response) for fast and slow Go trials. Averaging across Early and Late conditions, fast responses exhibited significantly steeper desynchronization slopes than slow responses. The relationship between desynchronization slope and response timing did not differ between the Early and Late conditions, indicating that temporal regularity shifted the timing of the underlying neural dynamics without altering their fundamental relationship to behavior

Decision times on Go trials were substantially faster in the Early than in the Late condition (471 vs. 742 ms from sphere disappearance, p < .001; Figure 1B), indicating that participants used learned temporal statistics of No-Go events to infer when responding became safe. The shift in decision times was accompanied by a corresponding temporal shift in alpha- and beta-band activity across fronto-central and parietal regions (Figure 1C). Before contact, alpha- and beta-band desynchronization was stronger in the Early condition, consistent with enhanced sensorimotor preparation (Figure 1D). These effects were maximal over parietal regions (Figure 1E), supporting the proposed role of the parietal cortex in translating temporal expectations into anticipatory action states^5,10^. We further quantified the slope of pre-response alpha- and beta-band desynchronization over parietal electrodes (8–40 Hz, −1.0 to −0.1 s). Across conditions, faster responses were preceded by a significantly steeper desynchronization slope than slower responses (t(19) = −3.71, p = .0015), with no interaction between response speed and temporal condition (all ps > .35; Figure 1F,G). Together, the contact- and response-locked analyses show that anticipatory oscillatory activity was both calibrated to learned temporal regularities and tightly linked to the timing of internally generated decisions.

## Discussion

A fundamental challenge for adaptive behavior is determining when to act in situations where no explicit signal indicates that action is required. Extending prevailing accounts of temporal expectation beyond stimulus–response behavior^2^, we demonstrate that temporal regularities enable the brain to infer when action becomes appropriate. Shift in decision timing due to No-Go temporal regularities was accompanied by corresponding changes in the dynamics of alpha- and beta-band desynchronization, consistent with previous evidence implicating parietal sensorimotor networks in temporal prediction and action preparation. Furthermore, the observation that steeper pre-response alpha- and beta-band desynchronization consistently preceded faster responses suggests that the rate at which sensorimotor activity evolves determines when an internally generated decision is translated into overt action. This is compatible with the idea that the rate of decrease of alpha-beta power acts as an internal clock, whose speed tunes to statistical regularities and whose variability directly drives behavioural variability. Our results identify a candidate neural mechanism through which temporal expectation is transformed into self-generated behavior, offering a framework for understanding how humans make decisions when the environment provides no direct instruction to act.

## Materials and Methods

### Participants

Twenty-five healthy volunteers participated in the study. Three participants were excluded from n behavioral analyses (i.e., reaction times) because of technical issues affecting response recording. All participants reported normal or corrected-to-normal vision and no history of neurological or psychiatric disorders. Written informed consent was obtained prior to participation. The study was part of a project approved by the Ethics Committee of the University of Milano-Bicocca (RM-772).

### Experimental Setup

The VR setup included a Meta Quest 2 head-mounted display (HMD) with a resolution of 1920 × 1832 pixels per eye. The HMD was connected via Oculus Link to an ASUS ROG Strix laptop equipped with an AMD Ryzen 9 5900HX CPU, 32 GB of RAM, and an NVIDIA GeForce RTX 3080 GPU, with a 17.3-inch display (1920 × 1080 resolution). The Temporal Inference Task was developed in virtual reality (VR) using Unity game engine. The virtual environment consisted of an empty room, where the participant’s arm and hand were dynamically mapped using the HMD’s built-in hand-tracking system. Tactile stimulations for No-Go trials were delivered using a 3 V coin-shaped vibrotactile actuator (30 mm diameter) attached to the palm of the participant’s right hand using medical tape. Vibrations lasted 100 ms. Stimulation onset was synchronized with events in the virtual environment using an Arduino Uno board that controlled the actuator, which was triggered by the VR software. The response button was held in the left hand.

### Temporal Inference Task

Participants performed a Temporal Inference Task in VR designed to investigate how temporal regularities guide action timing. Each trial consisted of a looming sequence in which a virtual sphere approached the participant’s hand (starting point 1.5 m, velocity 1.5 m/s). During most trials (80%), a brief tactile stimulus was delivered and signaled a No-Go trial. Participants were instructed to withhold their response whenever tactile stimulation was detected. On the remaining trials (20%), no tactile stimulation occurred, and participants were required to press a button as quickly as accurate as possible. Critically, Go trials were not associated with an explicit Go signal. Instead, participants had to infer when a response was appropriate from the absence of an expected tactile event. The timing of No-Go events differed between experimental conditions. Early condition: tactile stimulation occurred 300 ms before sphere–hand contact. Late condition: tactile stimulation occurred 200 ms after sphere disappearance. Each condition consisted of two blocks of 100 trials; each one lasted about 5 minutes.

### Procedure

Before the main experiment, participants completed two blocks to familiarize themselves with the Temporal Inference Task and the possible timings of No-Go events. Each familiarization block consisted of 100 trials. No-Go stimuli could occur at one of three time points relative to sphere– hand contact: Early (−300 ms), Sync (0 ms), or Late (+200 ms), each occurring on 30% of trials. The remaining 10% of trials were Go trials in which no tactile stimulation was delivered. Data from the familiarization phase were not included in the analyses reported here. Following familiarization, participants completed the Early and Late conditions. To counterbalance order effects, half of the participants performed the Early condition first, followed by a break and then the Late condition, whereas the remaining participants completed the conditions in the opposite order.

### Behavioral Analysis

Decision times were computed on Go trials relative to sphere–hand contact. Trial-by-trial reaction times were analyzed using linear mixed-effects models. Condition (Early vs. Late) was entered as a fixed effect and participant as a random intercept. Statistical analyses were performed in R.

### EEG Acquisition and Preprocessing

EEG was recorded from 64 scalp electrodes using an ActiCAP system (Brain Products) at a sampling rate of 1000 Hz. Data preprocessing was performed using Matlab toolbox EEGLab and involved bandpass filter (1–100 Hz), notch filtered at 50 Hz, and downsampled to 512 Hz. Artifact correction was performed using independent component analysis. Bad channels were interpolated using spherical interpolation. Data were subsequently re-referenced to the average reference. Epochs extended from −2 s to 2 s relative to sphere–hand contact.

### Time–Frequency Analysis

Time–frequency representations (TFRs) of oscillatory power were computed using the FieldTrip toolbox. Spectral decomposition was performed using complex Morlet wavelets with a fixed width of seven cycles, providing a balance between temporal and frequency resolution. Power estimates were calculated for frequencies ranging from 4 to 40 Hz in 1-Hz increments and for time points spanning −2 to 2 s relative to the event of interest, sampled at 10-ms intervals. To minimize edge artifacts, data were zero-padded to the maximum trial length prior to time–frequency decomposition. Baseline correction was performed using a pre-event interval from −2 to −1.5 s. Power values were converted to decibel (dB) units according to the formula 10×log10 (power/baseline), such that positive values indicate increases and negative values indicate decreases in power relative to baseline. Power estimates were then averaged across trials within each experimental condition to generate participant-specific TFRs, which were subsequently used to characterize oscillatory dynamics and served as the basis for all statistical analyses.

To assess differences in oscillatory power between experimental conditions, we used nonparametric cluster-based permutation tests implemented in FieldTrip. Analyses were restricted to electrodes spanning frontal, central, centro-parietal, parietal, and parieto-occipital scalp regions (F1, Fz, F2, F3, F4, F5, F6, FC1, FCz, FC2, FC3, FC4, C1, Cz, C2, C3, C4, CP1, CPz, CP2, CP3, CP4, P1, Pz, P2, P3, P4, P5, P6, PO3, POz, and PO4). Statistical testing was performed within a time window from −1 to 1.5 s relative to the moment of sphere–hand contact and across frequencies from 4 to 40 Hz. For each comparison, condition differences were assessed using a dependent-samples t-test with a Monte Carlo randomization procedure. Clusters were formed from adjacent time–frequency–channel samples exceeding the cluster-forming threshold, using the electrode neighborhood. A minimum of three neighboring channels was required for cluster formation.

Statistical significance was evaluated using a two-tailed cluster-level test, with a cluster-forming threshold of p < .025 and a cluster-level significance threshold of p < .025. Cluster-level statistics were computed as the sum of t-values within each cluster and evaluated against a permutation distribution (n = 500). This procedure allowed us to identify spatially, temporally, and spectrally extended patterns of oscillatory modulation associated with the experimental manipulation while controlling for multiple comparisons across the time–frequency space.

### Response-locked slope analysis

To assess whether the temporal evolution of oscillatory activity predicted response timing, we performed a response-locked analysis of alpha/beta power (8–40 Hz) averaged across parietal electrodes (Pz, P1, and P2). Single-trial time–frequency representations were first baseline-corrected relative to an event-locked pre-sphere interval (−1.5 to −1.0 s before sphere appearance) and were subsequently realigned to response onset. Within each condition, trials were split into fast and slow responses using a within-participant median reaction time. For each participant, the slope of pre-response desynchronization was estimated by fitting a first-order linear regression over the interval from −1.0 to −0.1 s before the response. Slope estimates were compared between fast and slow responses using paired-samples t-tests within the Early and Late conditions, as well as after averaging across conditions.

## Acknowledgments

This paper was supported by a grant funded by the Italian Ministry of University and Research (MUR) – PRIN 2022; Grant n. 2022-NAZ-0172 to Alberto Gallace.

## Data Availability statement

Experimental Data and Code to reproduce the analysis are available on OSF (https://osf.io/6tvm3/).

## References

1) Nobre, A. C. (2012). How can temporal expectations bias perception and action? In Attention and Time, 2010, 371–392

2) Nobre, A. C., & Van Ede, F. (2018). Anticipated moments: Temporal structure in attention. Nature Reviews Neuroscience, 19(1), 34–48

3) Correa, A., & Nobre, A. C. (2008). Neural modulation by regularity and passage of time. Journal of Neurophysiology, 100(3), 1649–1655

4) Cravo, A. M., Rohenkohl, G., Wyart, V., & Nobre, A. C. (2011). Endogenous modulation of low frequency oscillations by temporal expectations. Journal of neurophysiology, 106(6), 2964–2972.

5) Grabenhorst, M., Poeppel, D., & Michalareas, G. (2025). Neural signatures of temporal anticipation in human cortex represent event probability density. Nature Communications, 16(1), 2602

6) Rohenkohl, G., Cravo, A. M., Wyart, V., & Nobre, A. C. (2012). Temporal expectation improves the quality of sensory information. Journal of Neuroscience, 32(24), 8424–8428.

7) Rohenkohl, G., & Nobre, A. C. (2011). Alpha oscillations related to anticipatory attention follow temporal expectations. Journal of Neuroscience, 31(40), 14076–14084.

8) van Ede, F., Niklaus, M., & Nobre, A. C. (2017). Temporal expectations guide dynamic prioritization in visual working memory through attenuated α oscillations. Journal of Neuroscience, 37(2), 437–445.

9) Braga, A., & Schönwiesner, M. (2022). Neural Substrates and Models of Omission Responses and Predictive Processes. In Frontiers in Neural Circuits, 16, 799581.

10) Janssen, P., & Shadlen, M. N. (2005). A representation of the hazard rate of elapsed time in macaque area LIP. Nature Neuroscience, 8(2), 234–241.

